# Valve endothelial monolayer fissuring via RhoA activity induces 3D calcific aortic valve lesion emergence as revealed by a longitudinal live-imaging platform

**DOI:** 10.1101/2025.06.21.660489

**Authors:** Katherine Driscoll, Santosh Balakrishnan, G. Janani, Hrishika Gogineni, Alessandra Good, Andrea Huang, Nichaluk Leartprapun, Steven G Adie, Jonathan Butcher

## Abstract

Calcific aortic valve disease (CAVD) is a degenerative disease with wide prevalence in the aging population and a low survival rate after onset of symptoms, yet there are no effective pharmacological treatments. Many patients present to the clinic with symptoms at end-stages of CAVD, when the disease may be irreversible. The ability to identify and live-trace calcific lesion emergence in-vivo would allow for the identification of disease biomarkers and discovery of therapeutic targets at earlier, more treatable stages. In this work, we establish a new multimodal in-vitro CAVD platform consisting of lineage traced-VEC and VIC cells in a 3D model combined with live optical coherence and fluorescence microscopy to unravel VEC, VIC, and matrix transitional events during calcific lesion formation. We discover that fissuring of the endothelial monolayer combined with the formation of dense aggregates in adjacent regions is a key biomarker of the onset of lesion formation. This coincides with the formation of dense VIC and ECM conglomerates under endothelial aggregates, an additional biomarker of pathogenesis. Further, we discover that fibrotic tissue compaction is correlated with but not necessary for lesion formation. Additionally, we identify RhoA activation in disease-treated samples. We demonstrate that RhoA inhibition through ROCK, but not Rac1 inhibition, prevents delamination of the endothelial monolayer, fibrotic remodeling, and emergence of calcific lesions. Together, this work establishes a new longitudinal live-imaging platform that identifies emergent cell and matrix biological signatures of CAVD onset and enables the evaluation of therapeutic interventions.

**Graphical Abstract:** 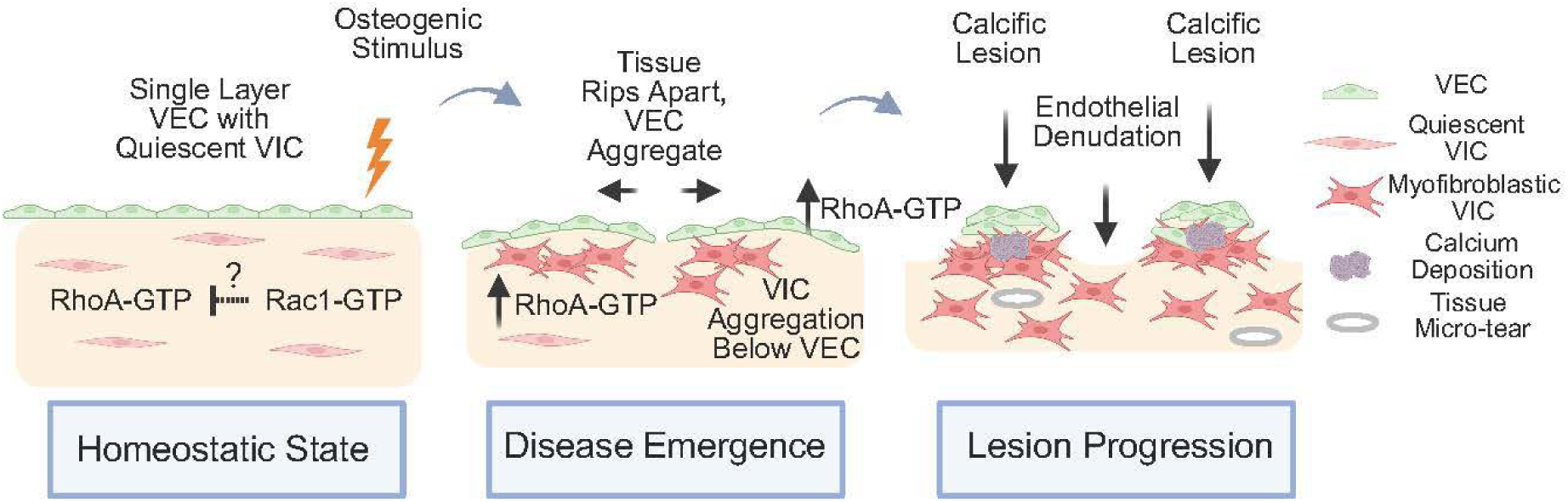

## Introduction

Calcific aortic valve disease (CAVD) is the gradual stiffening and calcification of the aortic valve in the heart, and if left untreated, eventually results in heart failure[1]. Aortic stenosis, a feature of early stage CAVD, affects over 12% of the population over age 75, and heart failure due to CAVD rose by 90% between 1990 and 2019[2,3]. However, there are no effective pharmacological therapeutics to date to treat the disease[4].

Healthy aortic valves have two major cell types: valvular endothelial cells (VEC) which line the valve and provide a barrier between the tissue and the blood, and valvular interstitial cells (VIC) which repair and maintain the integrity of the tissue. Calcific aortic valve disease presents with calcified nodules specifically on the fibrosa side of the leaflet, and fibrosis resulting in valve leaflet thickening and stiffening[1,5–8]. These changes reduce the compliance of the valvular leaflet, preventing adequate opening and closing over the cardiac cycle.

Although the presentation of CAVD is clear in its advanced stages, early spatiotemporal events leading to eventual CAVD pathogenesis are largely unknown[9,10]. CAVD patients often do not present to the clinic until they are symptomatic, at which point irreversible and lethal myocardial damage has often already taken place[11–14]. Identification of key pathogenic transition stages of the calcific disease process before the onset of stenosis would provide invaluable information for identifying CAVD patients in reversible stages of the disease. Identifying molecular mechanisms to target at these early and intermediate stages of the disease would also improve patient outcomes.

The new FDA Modernization Act 2.0 allows for complex in-vitro systems that accurately recapitulate pathogenesis to bypass animal testing requirements for clinical trials[15,16]. Most in-vitro studies in the CAVD field have utilized 2D VIC monoculture models of calcification, which do not recapitulate the multi-cellular complexity and three-dimensional nature of the disease[17–19]. More recent approaches have utilized 3D biomaterial approaches, but often only used one cell type (VIC). [17,20–23]. CAVD valve explants typically exhibit a breakdown in the VEC monolayer and basement membrane, which cannot be represented with a VIC-only experimental approach[6].

Our group has developed a tensioned co-culture model of CAVD consisting of a VIC-laden collagen gel lined with a VEC monolayer[24]. We previously discovered that in VEC+VIC co-culture, but not VEC-only or VIC-only monoculture, osteogenic-media treatment induces the formation of large raised 3D calcific lesions with evidence of collaborative participation of both cell types[24]. Additionally, these VIC-VEC lesions uniquely recapitulated spatial-patterning of molecular pathways in human valve calcifications, including osteoblastic VIC expression in calcific nodules, increased chondrogenic expression in the regions surrounding calcific nodules, and increased evidence of VEC EndMT in a multi-layer presentation[24].

Although we demonstrated the ability to faithfully induce spatiotemporally complex multicellular lesions in a high-throughput manner, the dynamics of the lesion formation process were unclear due to the lack of live-imaging. An understanding of the cascade of events leading to the formation of lesions would enable identification of signatures of early stage disease and elucidate the processes involved. This understanding is paramount, as such features have potential to be observable clinically and could be further monitored to guide treatment. The RhoA and Rac1 GTPase pathways are known to play a large role in cellular contractility and migration[25,26]. In particular, the RhoA pathway is known to induce myofibroblastic transformation, and fibrosis, a feature present in stenotic valves[27,28]. Evidence suggests that stenotic valves often, but not always, exhibit both fibrosis and calcification, although the relationship between the two is yet to be determined[28–30]. Our previous work, as well as others, have indicated that the RhoA and Rac1 pathways likely contribute to CAVD pathogenesis, although how these pathways affect 3D lesion emergence in real-time with both VIC and VEC cell types remains unknown[31–38].

To address the lack of time-course information of calcific lesion formation with previous studies, we utilized our previously described co-culture VEC-VIC model that resembles the 3D organization of the valve leaflet and produces lesions under osteogenic treatment[24]. The use of GFP-tagged VEC along with primary VIC allows VEC lineage tracing and to distinguish between VEC, VIC and the matrix within the model. A novel combination of 2D and 3D longitudinal imaging enables us to track changes at multiple scales over the culture period. End-point staining and IF imaging of the samples enables the quantification of calcium, EndMT-VEC, aSMA and other parameters of RhoA activity. This combination of model and imaging protocol yields a longitudinal, live-imaging platform that was used to study the CAVD process.

Bright-field and wide-field fluorescence were used for whole mount, longitudinal imaging, while a novel, home-built combined Optical Coherence Microscopy (OCM) and confocal microscopy system was used for high resolution imaging at both the surface and deep within the sample. This unique approach allowed us to observe the formation of calcific lesions in real-time, with the goal of identifying pivotal biological signatures of homeostatic breakdown in pathogenesis. Our findings provide a snapshot of key events in VEC, VIC, and matrix remodeling that occur before, during, and after lesion formation. We discovered that endothelial monolayer fissuring and aggregation with concurrent underlying VIC and ECM conglomerates are biomarkers of calcific pathogenesis, while fibrotic remodeling is correlated with but not necessary for lesion formation. After establishing key biomarkers of lesion formation with live-imaging, we evaluated the relationship between overall tissue compaction, as a proxy for fibrosis, to lesion formation. Then, we investigated whether the RhoA and Rac-1 GTPase pathways are active in lesion-forming samples. Finally, we used dynamic imaging to evaluate if and how therapeutic targets for RhoA and Rac1 might prevent pathogenesis differentially in VIC, VEC, and matrix (Figure 1).

**Figure 1.**
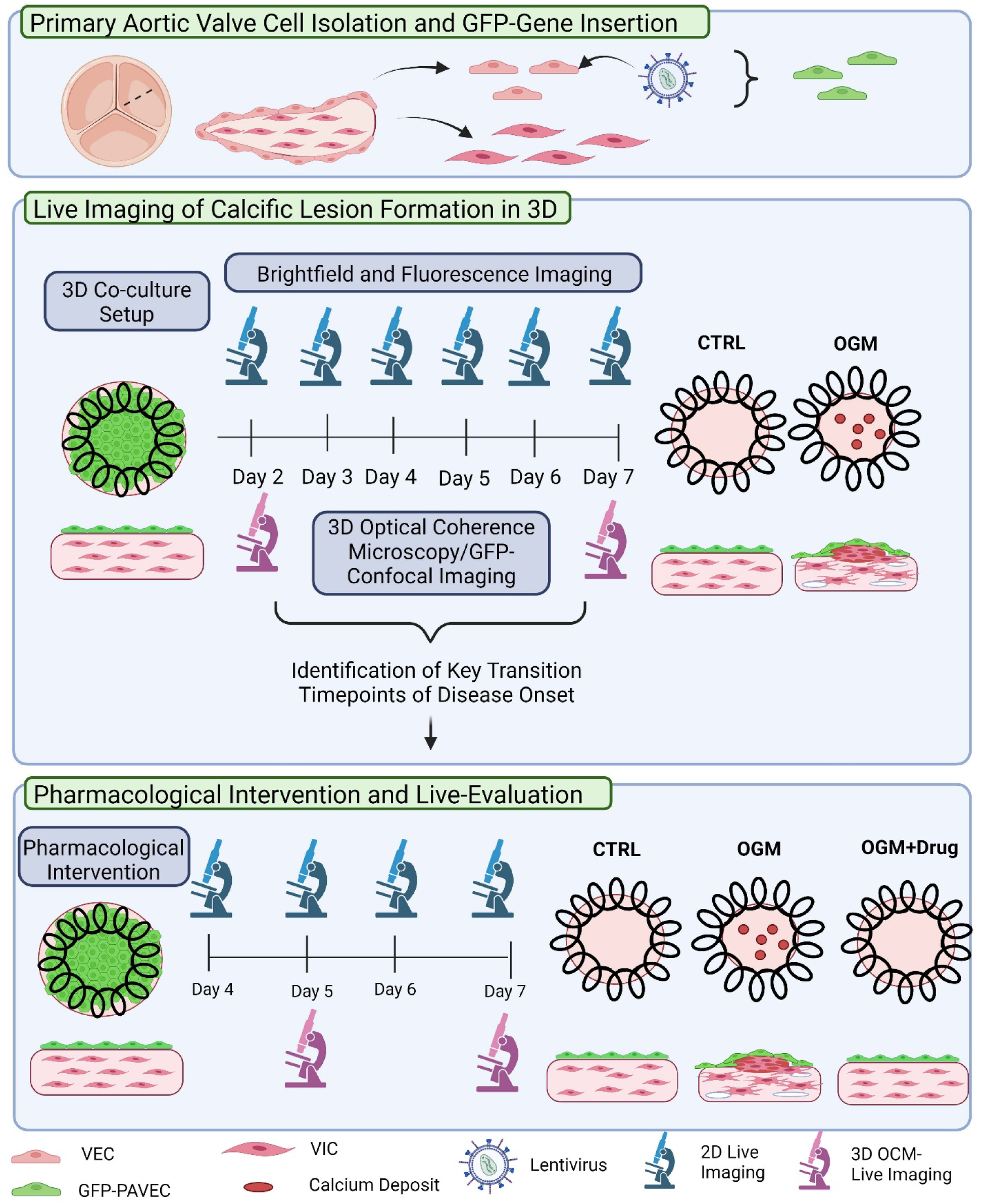
Study overview. Graphical representation of the methodology used for GFP-lineage tracing of GFP-VEC (top), live 2D and 3D timelapse imaging of calcific lesion formation (center), and live-evaluation of pharmacological intervention (bottom).

## Materials and Methods

### Porcine cell culture and cell isolation from aortic valve

Primary porcine VEC and VIC cells were isolated as previously described[24,39]. For routine cell culture prior to experimentation, porcine VIC were cultured in control media (DMEM + 10% FBS + 1x P/S) in tissue culture flasks, and were passaged at ∼50-75% confluency. Porcine VEC were cultured in tissue coated flasks coated with 50ug/mL rat-tail collagen type I (Enzo, Long Island, NY; IBIDI, Fitchburg, WI) in the same media as Porcine VIC with the addition of 50 U/mL heparin (VWR, Radnor, PA; Krackeler Scientific, Albany, NY), and were passaged at ∼80-95% confluency. Fresh porcine hearts were generously donated to us by Timberline Meats in Dundee, NY.

### Lentiviral GFP-transduction of primary Porcine endothelial cells

Ready-made lentivirus at a titer of 500,000 TU/mL containing GFP gene insert was purchased from Addgene (#17446-LV, Watertown, MA). Primary porcine VEC cells were cultured until confluent at passage 2. Polybrene media was prepared by adding 20µL of 10mg/mL stock (EMD Millipore, Burlington, MA) to 20mL DMEM+10%FBS, and in later iterations + heparin (50U/mL) without antibiotics. Lentivirus prep was prepared by adding Addgene 17446 lentivirus stock to polybrene media (5mL) at a 1:50 dilution per T75 flask. Endothelial cells were detached from the plate and 400,000 cells were added to 10mL polybrene+media base per T75 flask. 5mL virus suspension was added to the flask first, then 10mL VEC suspension for a total of 15mL/T75 flask. Virus+cells were incubated for 72 hours, then media was changed to DMEM+10% FBS+ 1xP/S + 50U/mL heparin. Cells were grown to confluency with frequent media changes upon observation of cell death and monitored for any abnormal changes in morphology. Upon confluency, cells were passaged at a 1:3 ratio.

GFP+ transfected VEC were purified using the BD melody FACS sorter. Sorting gates for collection included GFP+ cells and excluded DAPI+ cells to purify for viable GFP+ cells. The collected cells were centrifuged and the pellet was either plated following FACS sorting for further expansion or frozen immediately in 50% DMEM, 40% FBS, and 10% DMSO.

Transfected GFP+ VEC lots were validated utilizing brightfield microscopy for normal cobblestone morphology, fluorescence microscopy to determine %transfection efficiency, and qPCR to ensure continued expression of endothelial marker PECAM-1, and low expression of αSMA (Table 1).

**Table 1.**
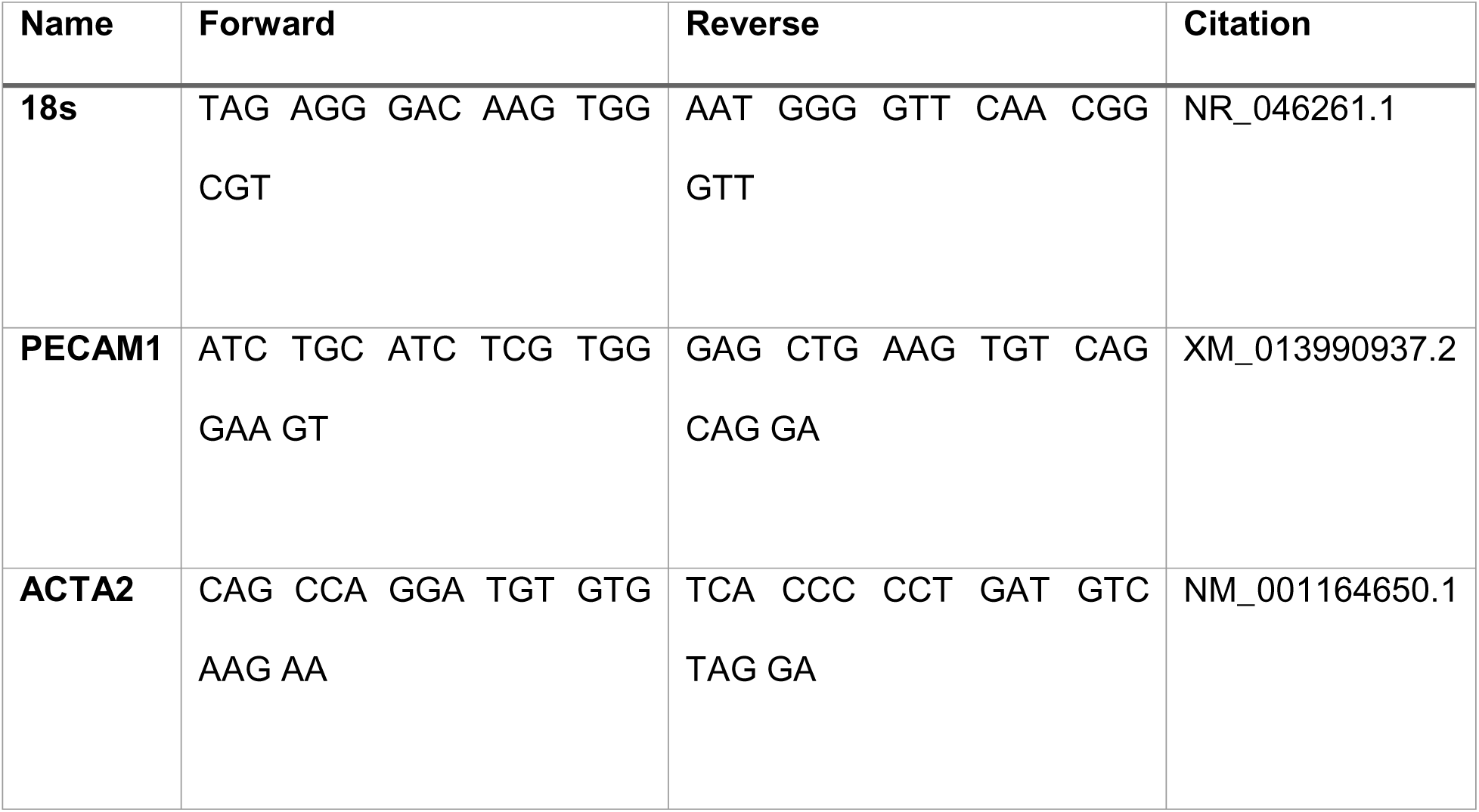
Forward and reverse primer sequences utilized for qPCR.

### 3D tissue engineered CAVD co-culture model fabrication and co-culture casting

The co-culture gel was cast inside of a circular cavity in a PDMS mold containing an embedded spring in the cavity as previously described[40]. One construct per well was inserted into a sterile 24-well plate prior to casting the co-culture gel. Next, porcine VIC cells were seeded in a 2mg/mL rat-tail collagen type I gel at a concentration of 400,000 cells/mL, with transfected GFP+ porcine VEC seeded on top at a concentration of 50,000 cells/cm^2.

The next day (day 1), media was changed to either control (CTRL) media (DMEM+10% FBS+ 1x P/S) or osteogenic media (OGM). OGM was prepared by adding 10mM B-glycerophosphate (EMD Milipore, Burlington, MA), 50µg/mL ascorbic acid (Sigma Aldrich, St. Louis, MO), and 100ng/mL dexamethasone (Sigma Aldrich, St. Louis, MO) to control media, as previously described[40]. Media was changed every other day of culture, and the experiment was terminated on day 7 of culture. Data from figures 2-6 was obtained through three independent experiments using 3 separate primary porcine VIC cell lines and 2 separate primary GFP-porcine VEC cell lines derived from 5 separate animals, unless otherwise noted.

**Figure 2.**
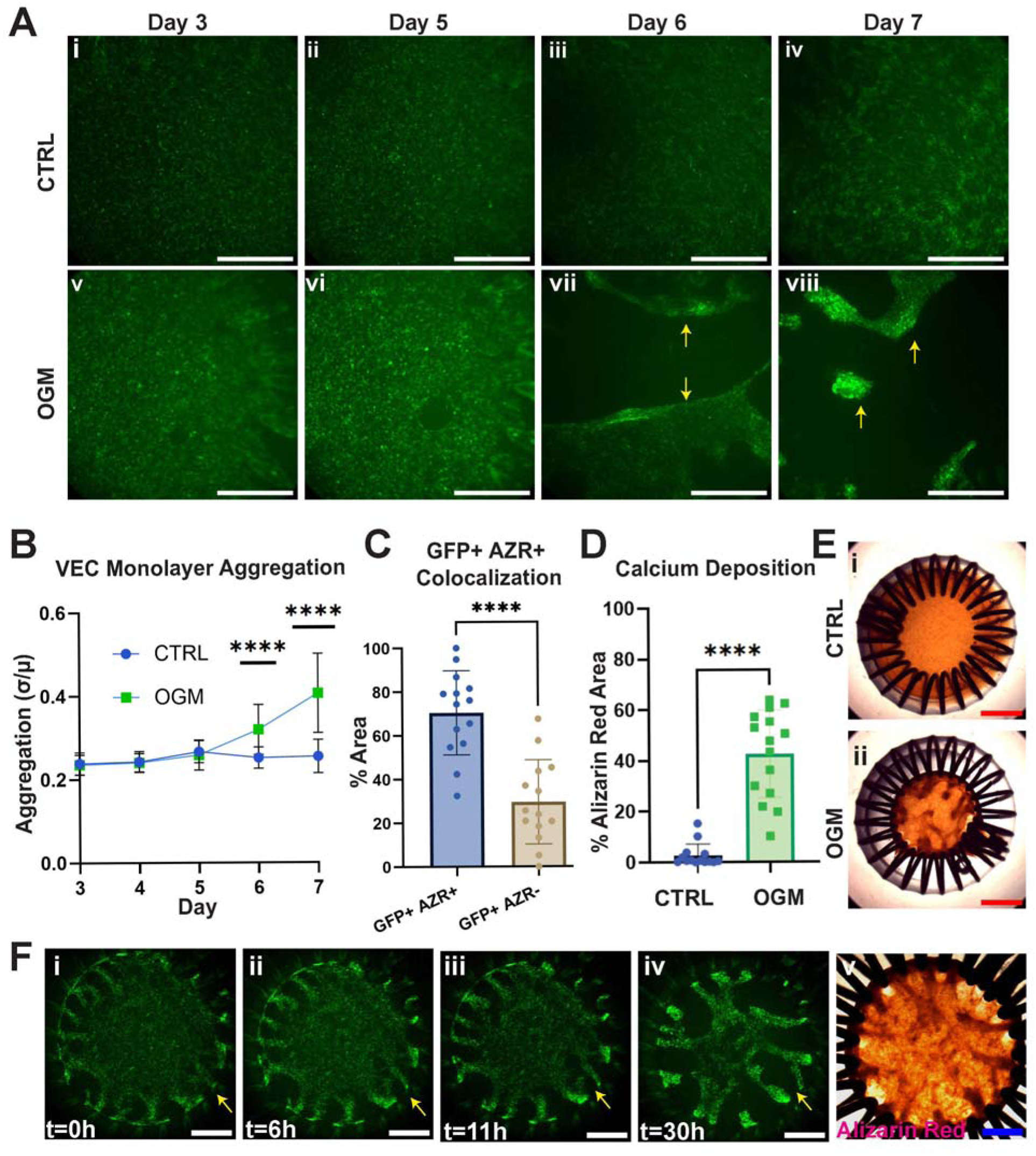
Daily live imaging of GFP+ VEC monolayer and corresponding fixed calcium stain in control and OGM-treated samples. A. Daily timelapse of the endothelial monolayer of control (i-iv) and osteogenic-media treated (v-viii) valve co-culture samples. Arrows point to regions of VEC clumping surrounded by empty space. Scalebar= 1mm. B: Daily GFP-VEC aggregation quantified (n=34-36). C: Quantification of percent overlapping GFP positive alizarin-red positive area normalized to GFP only area in OGM samples (n=14). D: Quantification of alizarin-red positive area in CTRL (Ei) and OGM (Eii) treated samples (n=14-15). E: Representative alizarin-red stain in CTRL and OGM samples, scalebar=2mm. F: (i-iv)Timelapse tracking GFP-VEC aggregation over 30h, at onset of aggregation formation. (v) Fixed alizarin-red stain in the same sample. Scalebars=1mm. . Student’s t-test was performed on control vs. OGM samples. Welch’s T-test was performed in cases of unequal variance.**** p ≤ 0.0001

### GFP-VEC surface live image acquisition and aggregation analysis

Constructs were imaged live daily from day 3 to day 7 with epifluorescence though the lid of the 24-well plate utilizing the Zeiss SteREO Discovery.V20 microscope. The same image acquisition parameters were used on all samples acquired on the same day. The imaging location was chosen with reference to the segment of the overlapping ends of the spring coil to ensure that the same region of each sample was imaged each day.

The amount of GFP-VEC aggregation was determined in Fiji/ImageJ[41] by the coefficient of variance of the pixel intensity in the green channel of each image. The coefficient of variance was calculated by dividing the standard deviation of pixel intensities by the mean image pixel intensity, where images with more heterogenous regions of extremely bright GFP+ aggregates and delaminated blank regions have higher standard deviations in proportion to the mean than images with cells in a uniform pattern. This analysis approach allowed the same quantification parameters to be applied to images with vastly different brightness/contrast due to aggregation in OGM and not CTRL samples, and images taken years apart with little batch-effect issues.

To calculate the normalized pixel intensity, the individual mean intensity value of each sample was normalized to the average intensity value of the imaging day across all samples. Three independent experiments were conducted with three different VIC and two different GFP-VEC primary cell lines from day 3 to day 7.

### Compaction image acquisition and analysis

Sample compaction was measured as a surrogate for fibrotic ECM remodeling[42]. Samples were imaged in brightfield daily using the Zeiss SteREO Discovery.V20 or a dissection microscope from day 2 to day 7 . Compaction analysis was performed in Fiji/ ImageJ by calculating the percent compaction for each CTRL to OGM sample across each day over 2-3 experiments, as previously described[24]. Briefly, percent compaction was calculated for each hydrogel by dividing the current gel area to the original area, determined by the area of the mold cavity (which is the initial gel area).

### Alizarin red staining and analysis

In order to quantify the amount of calcium deposition in each sample, we performed Alizarin red staining on day 7 with a 2% solution of Alizarin Red S (VWR, Radnor, PA) as previously described [40]. Between 0.5-1mL of 2% alizarin red stain was added to each sample for 20 minutes at room temperature with agitation, then the samples were rinsed 3x for >/= 15 minutes in 18mOhm water. The samples were left overnight to de-stain and the water was changed on the samples until it ran clear. Once the supernatant water was clear, samples were imaged in brightfield with the same imaging parameters applied across all images. Alizarin red images were acquired using brightfield microscopy on the Zeiss SteREO Discovery.V20 or a dissection microscope.

For alizarin red quantification, brightness, contrast, and color balance were corrected and a mask was applied to select only central area excluding the springs. Then, thresholding and gaussian blur were applied to select for the AZR+ regions. The percent alizarin red area was calculated by dividing the extracted AZR+ area by the mask of the selected region excluding the spring/non-sample area, and multiplying by 100. All image acquisition & processing settings for all samples were kept unchanged for each experiment across CTRL, OGM and/or drug treatment.

### Alizarin red and GFP-VEC co-localization analysis

To determine the relationship between endothelial delamination and calcification, we overlaid day 7 GFP-VEC surface images with images of the same samples fixed and stained with alizarin red for calcium. To align live-GFP images and fixed-alizarin red images taken at different times, images were registered in Adobe Photoshop using the spring as the fiducial. The AZR image was cropped to the same magnification and orientation of its corresponding GFP+ image and saved for further analysis in Fiji/ImageJ *(Supplementary Figure S2C)*.

The registered GFP and Alizarin images were converted to masks in Fiji/ ImageJ by applying thresholding and Gaussian blur. The intersection of the segmented GFP+ and AZR+ masks was calculated using the image calculator logical “AND” in Fiji/ImageJ, and the area was measured *(Supplementary Figure S2C).* The area of the GFP-only and the AZR-only masks was also calculated. For comparison to alizarin-red only area, the AZR+ GFP+ area was divided by the total AZR+ area to get the %AZR+ GFP+ area, and the %AZR+ GFP-area was calculated by subtracting the %AZR+ GFP+ area from 100%. The %AZR+ GFP+ area normalized to GFP+ area was calculated in the same fashion.

### Combined Optical Coherence Microscopy and Confocal imaging

To enable a more nuanced understanding of the live-cellular and matrix changes that occurred in 3D to form calcific lesions, we employed a novel imaging approach that combines GFP-confocal microscopy with label-free 3D optical coherence microscopy (OCM) (Figure 1 center). We performed this multimodal imaging technique in the same control and OGM-treated samples to represent timepoints before (day 2) and after (day 7) the onset of CAVD. For OCM, light from a broadband superluminescent diode source, with a center wavelength of ∼1300nm, bandwidth of ∼175nm (EBS300061-04, Exalos AG) was split into a reference and sample arm using a 50:50 fiber coupler. The interference between the light returning from the two arms was detected using a spectrometer (Cobra 1300, Wasatch Photonics Inc.). The confocal microscopy sub-system employed a 488nm excitation laser and two photodetectors (HC125-02, Hamamatsu corp.), one that detected reflected light (483-493nm) and the other detecting fluorescence emission with wavelengths 505-545nm. Both OCM and confocal sub-systems shared a common optical path to the sample. Samples were imaged in the inverted configuration, using a 20x, (0.4 effective NA) objective (LC Plan N/NIR 20x, Olympus Inc.). Samples were placed in a CO2 microscope stage incubator (Uno-Plus, Okolab) that was held on a motorized platform (ZST225B, Thorlabs Inc.) for live imaging. The lateral resolution for the OCM and confocal imaging was 1.6 μm and 0.7 μm respectively, while the axial resolution was determined to be 7.5 μm (in air) and 5.6 μm respectively. OCM volumes were obtained at a single focal plane position and then computationally refocused[43]. The image intensity was normalized in depth and gamma compressed for ease of visualization. Confocal stacks were acquired with a 1 x 1mm field of view (FOV) and 1024 x 1024 pixels in X, Y with a Z spacing of 10 μm. The reconstructed OCM volumes were 1x 1 x ∼2.6 mm in X, Y and Z, with 1024 pixels in each dimension. Automated acquisition and saving of data was accomplished using a custom-built program (LabVIEW, National Instruments Inc.).

Maximum intensity projections (MIP) of the confocal fluorescence stacks were obtained using Fiji/ImageJ and were used to visualize the VEC distribution. These MIP images were analyzed using CellProfiler[44] to obtain VEC area and eccentricity. Representative results of the VEC segmentation are shown in Supplementary Figure S3.

Multiple OCM volumes were acquired in quick succession and a standard deviation projection was used to reduce the contribution of the extracellular matrix and to emphasize VIC bodies. Fiji/ImageJ was used to produce two OCM XY MIP frames: one MIP (“Surface”) at the surface that included all planes from the sample surface (the top and bottom axial indices were manually identified based on the appearance of the surface) and another MIP (“Interior”) over a ∼20 μm thick section, located ∼100 μm below the surface. The VIC bodies on the interior MIP were segmented using iLastik[45] and their area coverage and width were quantified using CellProfiler by using the segmentation probabilities as the input. Additional details on the segmentation, analysis procedure as well as intermediate results and validation are shown in Supplementary Figure S3.

### Whole-mount and cross-sectional immunofluorescent staining

Whole mount immunofluorescent staining was performed as previously described[40]. Briefly, samples were fixed in ∼4% PFA overnight, then permeabilized and blocked. Primary GFP-antibody was added overnight (see Table 2), and secondary antibody was added the next day for 1 hour at a 1:500 dilution at the same time as phalloidin-555 at a 1:1000 dilution. DAPI counterstain was performed and samples were imaged using the Zeiss LSM 880 with a 20x water immersion objective.

**Table 2.** List of primary antibodies and corresponding concentrations used for immunofluorescent staining.

| **Antibody** | **Manufacturer** | **Catalog #** | **Dilution/Concentration** |
| --- | --- | --- | --- |
| Chicken anti-GFP | Invitrogen | A10262 | 1:200 |
| Rabbit anti- 1-smooth- | Abcam | Ab5694 | 1:100 |
| muscle actin |  |  |  |
| Mouse anti-RhoA-GTP | New East Biosciences | 26904 | 1:100 |
| Mouse anti-Rac1-GTP | New East Biosciences | 26903 | 1:100 |

Cross-sectional immunofluorescent staining was also performed as previously described[40]. Briefly, co-culture constructs were fixed in 4% (v/v) PFA and processed for paraffin sectioning. 10µm sections were rehydrated and subjected to heat-mediated antigen retrieval using citrate buffer (Krackeler Scientific, Albany, NY) at 60°C overnight. Following antigen retrieval, sections were permeabilized and blocked with 5% (v/v) goat serum. The sections were washed and incubated for 1 hour at room temperature or at 4°C overnight in primary antibody solution prepared at specific dilutions listed in Table 2. The next day, samples were rinsed and secondary antibody was added at a 1:500 dilution for 1 hour. After rinsing and autofluorescence quenching with Vectorlabs TrueVIEW® Autofluorescence Quenching kit (Vectorlabs, Newark, CA), samples were incubated with DAPI, rinsed, and coverslipped with mounting media. Samples were imaged using a Zeiss LSM 710 confocal microscope with a 25x water immersion objective. Images were processed using Fiji/ImageJ and the percentage of positive cells for each marker was calculated for further analysis.

### RhoA-family pharmacological modulation and analysis

For RhoA-family pharmacological modulation, Rac-1 inhibitor NSC23766 (Sigma Aldrich, St. Louis, MO) and ROCK inhibitor Y-27632 (Sigma Aldrich, St. Louis, MO) were added at a 10µM working concentration to OGM-media on day 1 of treatment[31,46]. Thrombin (Sigma Aldrich, St. Louis, MO) was added at a 1 U/mL working concentration on day 1 to control and OGM-media for activation of Rho-A[47]. Drug treatments were refreshed every other day upon each media change.

Two independent experiments encompassing two GFP-porcine VEC primary cell lines and two porcine VIC cell lines from four separate animals were performed. Daily compaction and GFP-VEC live imaging were performed each day from day 4 to day 7. Combined OCM & Confocal timelapse imaging was performed on the same sample on day 5 and day 7. At day 7, the samples were Alizarin-red stained and imaged. Image acquisition and analysis for these steps were performed as previously described.

### Statistical analysis

All statistics were performed using GraphPad Prism 10. Only p-values less than or equal to 0.05 are represented on graphs and considered significant. Data on all graphs depicts the mean and standard deviation unless otherwise specified. The specific statistical test performed for each graph is denoted in the corresponding figure caption. The individual datapoint spread for all graphs are portrayed either in the main-text or supplementary figures. Only biological replicates were used as individual datapoints for analysis. The sample number for each figure is displayed in the corresponding figure caption as “n=” This work was performed in accordance with all state and federal laws, under Cornell IBC MUA# 15958

## Results

### Live imaging over 7 days reveals rapid delamination and aggregation of endothelial monolayer under osteogenic conditions

We utilized lentivirus transfection to create GFP+ primary valvular endothelial cell lines for live tracking of the valvular endothelium in our 3D VIC+VEC co-culture system *(Figure 1 (top), Supplementary figure S1 A-E).* The same locations of CTRL and OGM treated samples were live-imaged every day from day 3 to day 7 of culture (*Figure 1 (center))*. Over the course of the culture period, the endothelium of control samples remained homogenous, with some uniform cell loss in the endothelial monolayer occurring by day 7 *(Figure 2 A*). In contrast, the endothelium of the osteogenic-treated samples were confluent until day 6 of culture. At this time, massive fissures emerged in the endothelium *(Figure 2 A vii, yellow arrows*). These endothelial fissures rapidly propagated leaving behind large areas of delamination by day 7 *(Figure 2 A viii)*. The endothelial cells that formerly comprised the monolayer were densely clumped together adjacent to delaminated regions (*Figure 2 A viii).* This process of endothelial aggregation was quantified by measuring the monolayer coefficient of variance, and the VEC monolayer of OGM samples was statistically more heterogenous than control samples beginning at day 6, and this increased further through day 7. *(Figure 2 B)*.

To rule out endothelial death as a cause of delamination, we quantified the average signal intensity of OGM-treated samples compared to control samples *(supplementary figure S1 F-G).* Despite the presence of large regions devoid of endothelium, the OGM-treated samples exhibited a higher average intensity compared to control samples on day 7, indicating that endothelial death was not likely the primary mechanism of monolayer delamination in OGM samples *(supplementary figure S1 F-G)*.

To capture this VEC aggregation and delamination process in more detail, we live-imaged the endothelium of one sample starting at day 6 at low magnification over the entire culture area at 0, +6h, +11h, and +30 hours. *(Figure 2 F*). At Day 6 t=0, the endothelial monolayer demonstrated early signs of aggregation near the edges of the sample. Between regions of endothelial aggregations, VEC cells with an abnormally elongated morphology were observed to be migrating or being forcibly separated (Figure 2F i, yellow arrow). By t=30h, few elongated cells were present and a pattern of dense VEC clumps surrounded by large, delaminated regions was apparent (Figure 2F iv, yellow arrow).

### Large calcific lesions are colocalized with regions of endothelial aggregations

We fixed and alizarin-red calcium stained the same samples that were live GFP-imaged *(Figure 2D).* As expected, we saw significantly more calcium deposition in OGM vs. CTRL samples *(Figure 2E).* The colocalization of calcium deposits and VEC aggregation in OGM-samples can be seen on Figure 2F iv and v, which represent the VEC-GFP and the Alizarin-red stained images for the same sample. Quantification of the area of co-localization revealed that there was significantly higher VEC + calcium colocalized area than VEC alone or calcification alone *(Figure 2 C, Supplementary Figure S2).* The area surrounding regions of calcific lesions/VEC clumps was often devoid of endothelial cells.

### Combined OCM and confocal imaging reveals timelapse imaging reveals divergent pathogenesis at the surface vs deeper in the tissue in osteogenic-treated samples

The same CTRL and OGM samples were imaged with the combined OCM, Confocal system on day 2 and day 7, and the results are shown in Figure 3. The observations closely match those obtained from the 2D imaging described in the previous section. On day 2, there were no discernable differences between control and OGM-treated samples in the GFP-VEC monolayer *(Figure 3A),* surface collagen matrix/basal lamina *(Figure 3B),* deep collagen matrix *(Figure 3C, 3D)* and VIC cell morphology, *(Figure 3C, 3F)*, except for an increase in VEC cell area in OGM-treated samples *(Figure 3E).* The GFP-VEC in both groups had a very classical rounded morphology with equal eccentricity *(Figure 3A, 3E),* and the VIC cells were homogenously distributed with a thin, elongated morphology (Figure 3C).

**Figure 3.**
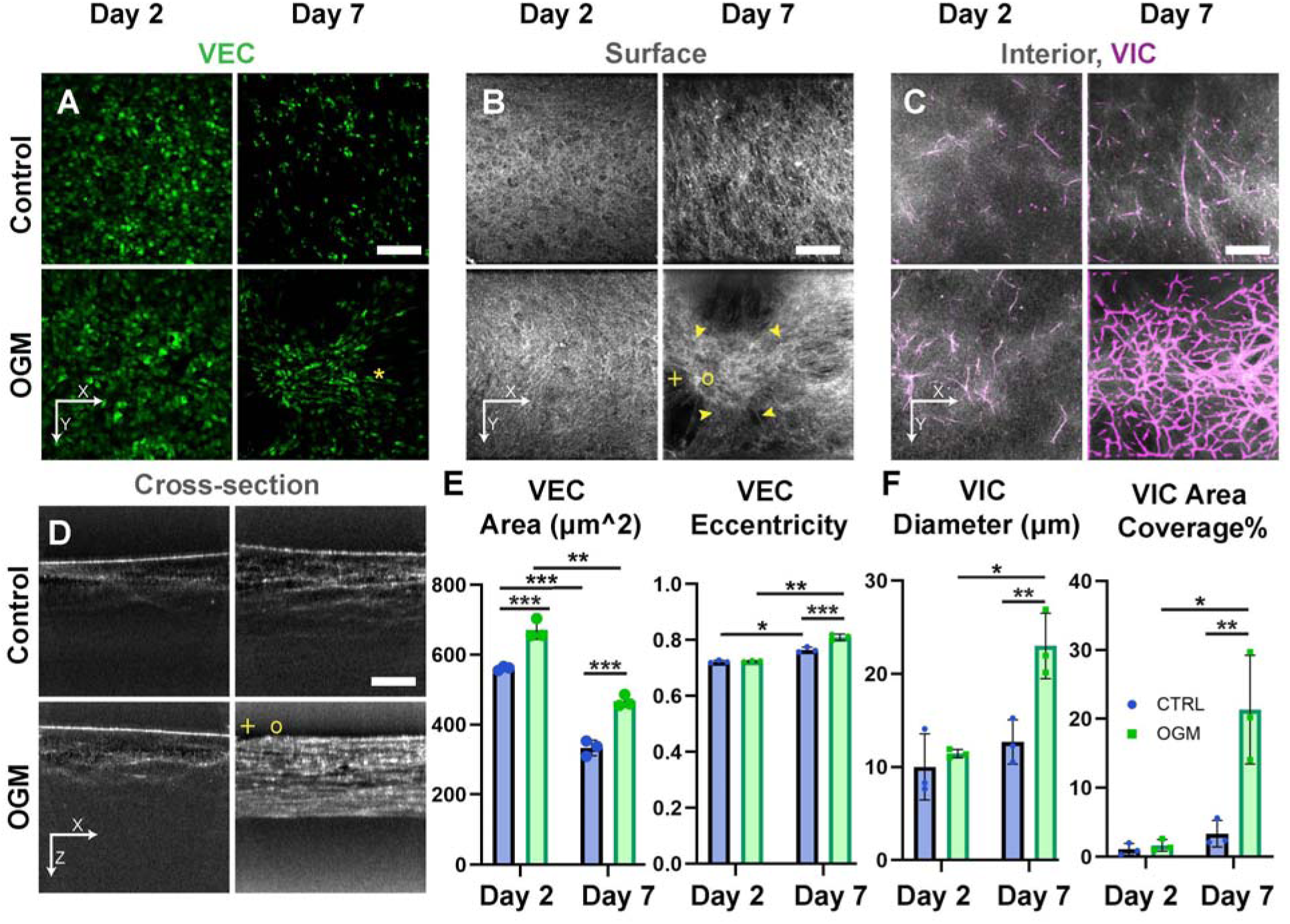
Comparison of the same control and OGM samples on Day 2 and Day 7. A: VEC confocal fluorescence MIP images showing differences in VEC morphology and migration. ‘*’ marks elongated VECs. B: OCM maximum intensity projection of images at the surface show differences in tissue surface remodeling. The region with a lesion is marked by ‘o’ and arrows indicate regions with highly aligned structures around the lesion, ‘+’ marks the region with a surface tear. C: OCM maximum intensity projection of images of a ∼ 20 µm thick section, ∼ 100 µm below the surface showing thicker VICs with significantly greater density in the OGM case. Computationally segmented VIC cells are shown overlaid in magenta. D: OCM cross-sectional images. ‘o’ and ‘+’ denote corresponding features shown in part B. E: VEC morphological properties (n=3). F. VIC diameter and percentage area coverage (n=3). Scale bars are 200 µm. Two-way ANOVA performed with Bonferroni’s multiple comparisons test. * p≤0.05, **p≤0.01, ***p≤0.001

However, by day 7, dramatic differences in the GFP-VEC monolayer, surface tissue, deep matrix, and VIC were observed in OGM samples compared to day 2 *(Figure 3 A-D, left vs. right columns).* We observed aggregation and delamination of the endothelial monolayer similarly to our observations utilizing 2D fluorescent imaging *(Figure 3 A).* In day 7 OGM-treated samples, the endothelial cell area decreased, and the VEC eccentricity increased compared to OGM-treated day 2 samples and compared to control day 7 samples. *(Figure 3A ‘*’ marker and 3E).* With OCM imaging, we visualized the tissue directly underneath the disrupted endothelium and identified a highly scattering dense “epicenter” of entangled VIC+matrix *(Figure 3 B,D circle).* The tissue surrounding this epicenter had a radially-aligned pattern that was more dense and organized than the basal lamina in OGM-treated samples on day 2 *(Figure 3 B, arrows)*. Tears in the basal lamina were often observed surrounding the aggregated lesion space *(Figure 3 B, D, plus sign)*. Deeper in the tissue, we observed a uniform lattice-like “honeycomb” network patterning of the VIC-cells and matrix *(Figure 3 C).* The VIC cells had a larger area coverage than day 2, and also had a much thicker morphology *(Figure 3 F).* Micro-tearing in the deep collagen matrix was also observed on day 7 in the OGM sample as evidenced by the dark pockets seen on the cross-sectional image in Figure 3D. Comparing Figure 3B with 3C for the day 7 OGM, it can be seen that the VIC and collagen matrix network pattern below the surface of the tissue are visually distinct from the shape of the lesions on the surface of the sample.

In contrast, in control samples on day 7, cell size of the GFP-VECs was reduced significantly, and the eccentricity was only slightly increased compared to day 2 *(Figure 3A, 3E)* and there were no differences observed in the VIC morphology *(Figure 3 F).* There was also some visual evidence of some minor collagen reorganization in control samples on day 7 evidenced by some bright matrix aggregations in the OCM images that differed from the homogeneously distributed collagen fibrils present on day 2 *(Figure 3C, Control day 2 vs day 7)*.

### OGM sample 24-hour timelapse depicts endothelium and underlying matrix+interstitial cell conglomerate concurrently “ripping apart” to form calcific lesion

We sought to distinguish whether the VEC delamination and aggregation process preceded or followed underlying VIC and matrix tearing during lesion formation. To do this, we performed a 24-hour timelapse during nodule formation on an OGM sample using the OCM/GFP-confocal live imaging system. The results depicting the endothelial monolayer, surface VIC/basal lamina, and deep VIC+collagen matrix during lesion formation are shown in Figure 4. The endothelium *(Figure 4A*) and basal lamina/VIC conglomerates *(Figure 4B*) fissured and formed aggregations simultaneously, and it was not obviously apparent that one process was preceded or followed the other. We captured further evidence of endothelial cells with abnormal elongated cell morphology separating in a similar fashion to our 2D widefield images *(Figure 4,A)*.

**Figure 4.**
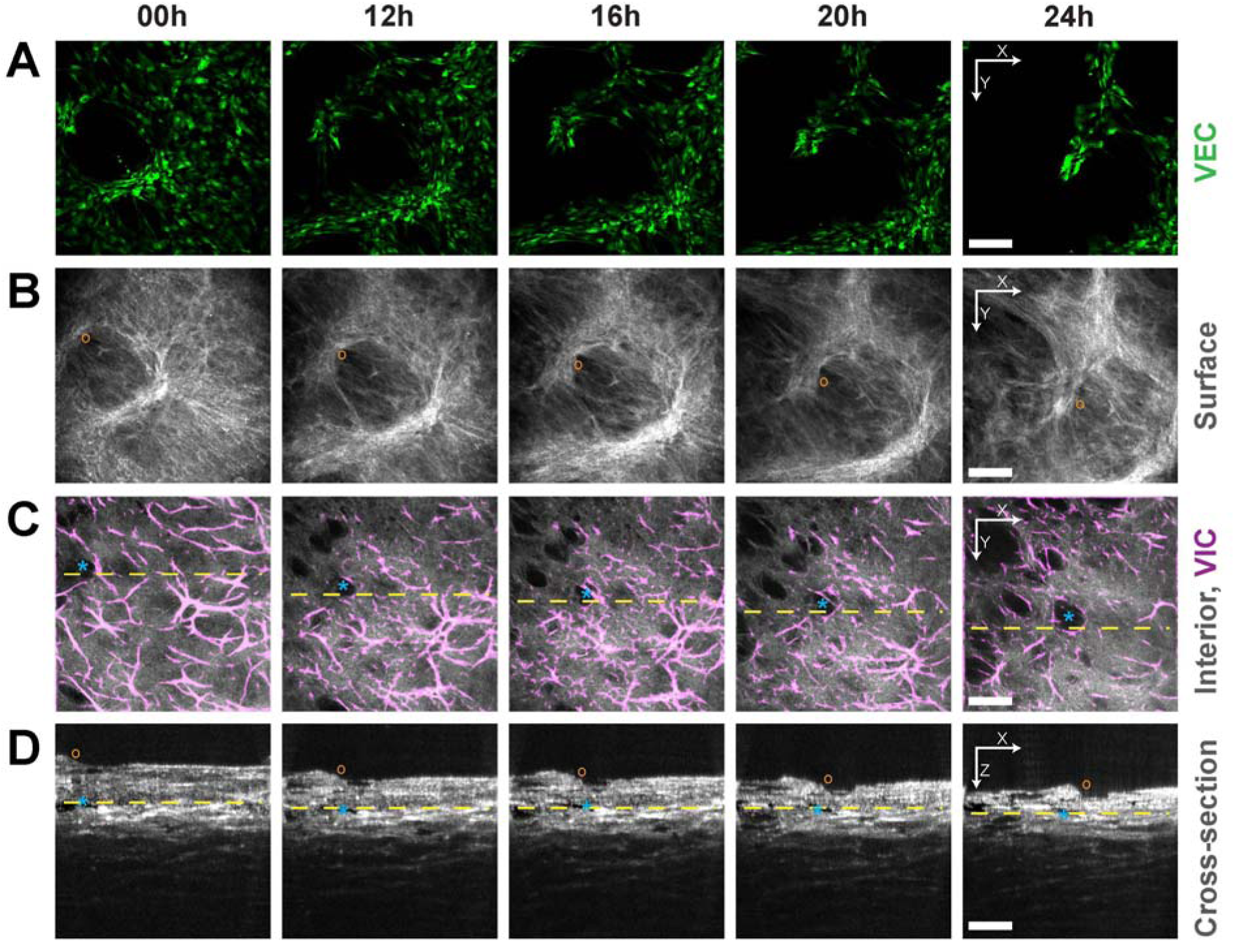
Live imaging of lesion formation on an OGM sample from Day 5 to 6. A: VEC confocal fluorescence MIP images showing the denudation of the VEC monolayer. B: Confocal reflectance MIP images depicting a lesion forming at the surface. C: OCM MIP images of a ∼ 20 µm thick section, ∼ 100 µm below the surface showing the propagation of micro-tears within the sample. Computationally segmented VIC cells are shown overlaid in magenta. D: OCM cross-sectional images illustrate the sample compaction and remodeling in depth. The ‘o’ markers in B, D track the lesion and ‘*’ markers in C, D label a micro-tear seen over time on the different images. The dashed lines in C indicate the Y position at which the XZ images in D are shown. The dashed lines in D denote the depth at which the projections in C were obtained. Scale bars are 200 µm.

Deep within the tissue, we captured microtear formation and propagation *(Figure 4C, 4D).* Small microtears emerged at the beginning of the imaging session in the top left corner of the FOV (denoted by ‘*’) and grew substantially in size over the 24-hour timelapse. Similar to the earlier results, the VIC honeycomb structure *(Figure 4C)* appeared to be independent of the aggregation and lesion formation processes at the surface of the sample *(Figure 4B)*.

Figure 4D shows cross-sectional views of the sample in depth, and the same features from Figure 4B and 4C can be seen sideways. Two aggregated features are seen on Figure 4B, pulling the surface apart and leaving behind a void between them as denoted by the ‘o’. Several micro-tears deep within the tissue that were not spatially arranged in alignment with the surface were also noted (*Figure 4C, 4D ‘*’*)

### Immunofluorescent Staining Depicts Dense VIC Bundles Underneath VEC Aggregations

To gain a better understanding of the 3D spatial orientation of VIC cells in relation to the VEC cell-aggregates, we performed fixed whole-mount staining and confocal imaging of OGM samples for VIC (f-actin) and VEC(GFP) at day 7 of culture, and the representative results are presented in Figure 5A. There were dense aggregations of interstitial cells directly underlying the regions of clumped endothelial cells, and not in regions that were devoid of endothelium (Figure 5 *A ii)*. Below and surrounding these dense aggregations were VIC that had a more classical interstitial cell morphology.

**Figure 5.**
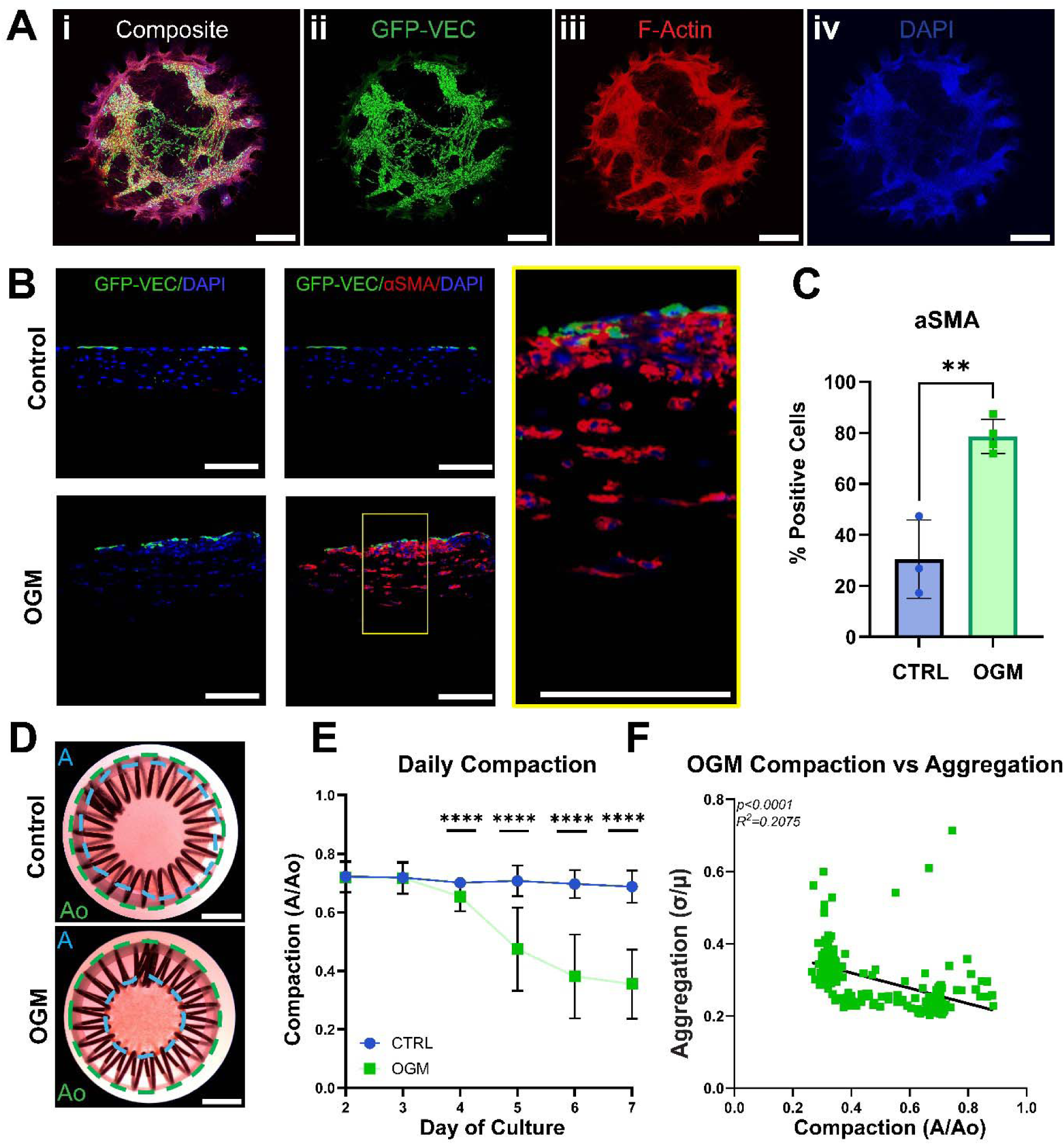
Immunofluorescent staining for GFP-VEC and activated VIC, and daily live bulk-compaction of control and OGM-treated samples. A. Whole-mount confocal immunofluorescence image of OGM-sample stained for GFP-VEC, F-Actin, and DAPI. Scalebar= 1mm. B: Cross-sectional images of CTRL and OGM-treated samples stained with GFP(green), aSMA(red), and DAPI(blue) Scalebars=100µm. C: Percentage positive aSMA cells in control and OGM-treated cells analyzed with student’s t-test (n=3-4). D: Brightfield images of CTRL and OGM samples demonstrating compaction measurement. Scalebars=2mm. E: Daily compaction summary of CTRL and OGM-treated samples (n=38-44). Welch’s t-test performed on CTRL vs OGM. F: Linear regression correlation analysis of GFP-VEC aggregation and compaction in OGM-treated samples day3 -day7 (n=166). ** p≤0.01, **** p≤0.0001

**Figure 6.**
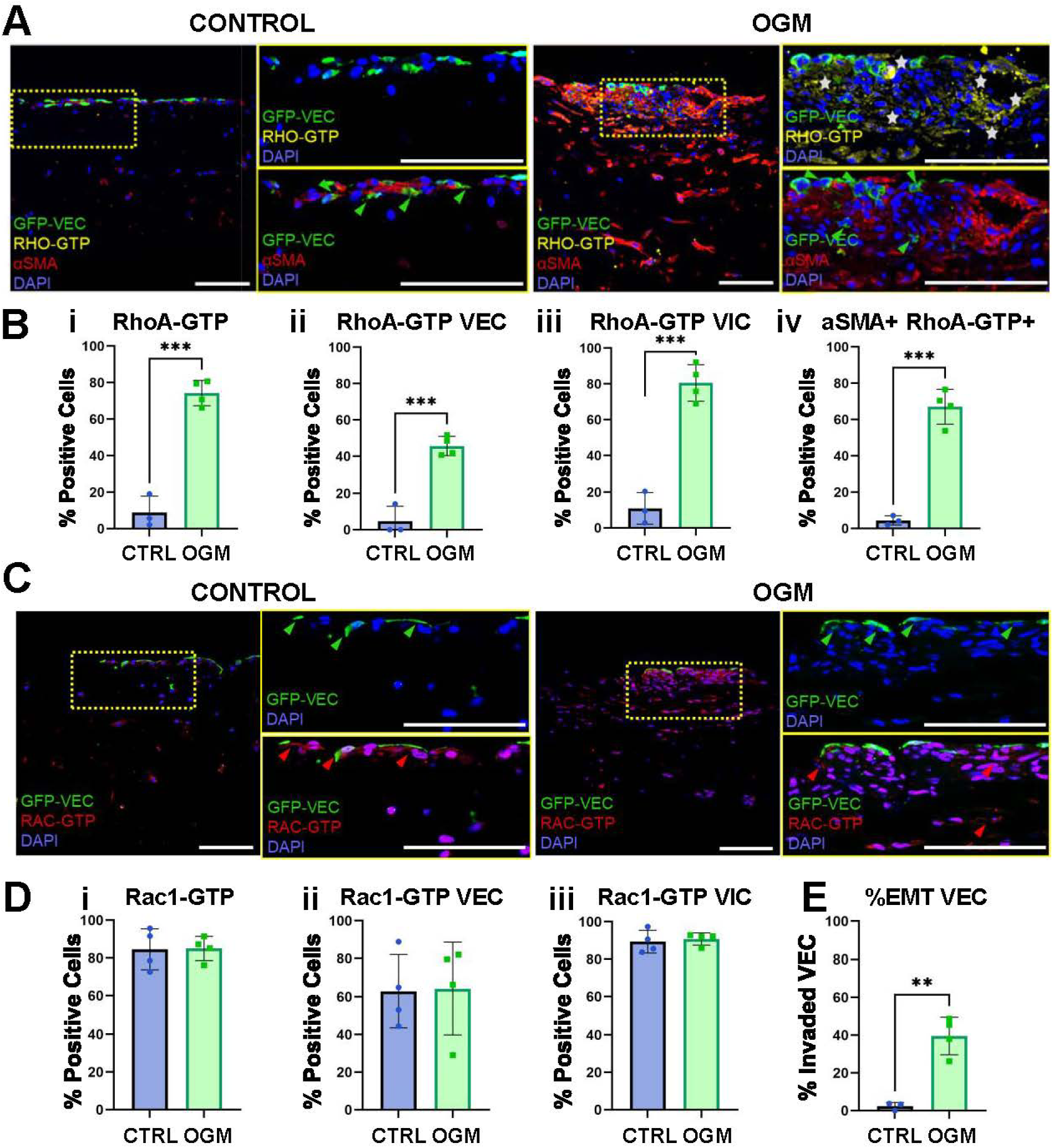
Immunofluorescence staining and quantification of Rho-GTP, aSMA, Rac-GTP, and EMT-VEC in control and OGM co-culture cross-sections. A. Cross-sectional images of Control and OGM samples stained with GFP-VEC (green), RhoA-GTP (yellow), aSMA (red), and DAPI (blue). Scalebars = 100 µm. Green arrows and white stars represent GFP-VEC and RhoA-GTP cells respectively. B. Percentage positive cells for (i) Rho-GTP, (ii) Rho-GTP expressed in VEC, (iii) RhoA-GTP expressed in VIC, and (iv) RhoA-GTP aSMA double stain in CTRL and OGM samples analyzed with student’s t-test(n=3-4). C. Cross-sectional images of Control and OGM samples stained with GFP-VEC (green), Rac-GTP (yellow), and DAPI (blue). Scalebars = 100 µm. Green arrows and red arrows represent GFP-VEC and Rac-GTP cells respectively. D. Percentage positive cells for (i) Rac-GTP, (ii) Rac-GTP expressed in VEC, and (iii) Rac-GTP expressed in VIC in CTRL and OGM samples analyzed with student’s t-test (n=4). E. Percentage positive cells for invading EMT VEC in CTRL and OGM samples analyzed with student’s t-test.** p≤0.01, *** p≤0.001.

We performed additional immunofluorescent staining on sections of control and OGM-treated samples to generate a cross-sectional view (Figure 5 *B).* In control samples, we typically saw a thin, single-layer endothelium. In contrast, OGM-treated samples often had clumped endothelium in multiple layers, and quantification of this feature revealed that there were significantly more endothelial cells below the surface of the sample in OGM-treated samples compared to control (Figure 6 *E)* Immunofluorescent staining for αSMA, a myofibroblastic marker, revealed a dramatic increase in αSMA expression in OGM vs control samples, especially in the surface-lesion space (Figure 5 *B, C)*.

### Macroscopic ECM compaction is correlated with but not necessary for VEC delamination process

We live-imaged control and OGM-treated samples daily and quantified the amount of compaction as a measure of fibrotic ECM remodeling (Figure 5 *D, E).* On day 4, OGM-samples became significantly more compacted than control samples, and this difference increased through day 7 (Figure 5 *E)*.

To discern whether there was a correlation between ECM compaction and VEC aggregation, we plotted relative aggregation vs. compaction for all OGM samples from day 3 to day 7 (Figure 5 *F).* Linear regression analysis revealed a weak but significant correlation between compaction and VEC aggregation. However, there were samples that had high levels of VEC aggregation with lower levels of compaction, indicating that VEC aggregation was not dependent on bulk-tissue compaction.

### Cell type specific GTPase activity associates with calcified lesion formation

There was significantly more RhoA-GTP activation in all OGM-cells and in both OGM-VEC and VIC compared to control, with an average of > 40% of VEC and > 70% of VIC expressing RhoA-GTP (Figure 6A*, 6B i,ii, iii ).* There was also abundant expression of Rac1-GTP in OGM-treated VEC and VIC, with an average of >60% of VEC and 80% of VIC expressing Rac1-GTP (Figure 6C*, 6D ii, iii).* However, this expression was not statistically different from the control samples for each cell type or for all cells (Figure 6D *i,ii,iii).* Quantification of the co-localization of cells expressing RhoA-GTP and aSMA revealed that greater than ∼60% of all cells in osteogenic-treated samples exhibited co-expression of both proteins, in contrast to control-treated samples, which had fewer than 10% of cells with co-expression (Figure 6 *B iv)*.

### RhoA, but not Rac1 inhibition prevents endothelial delamination, calcific lesion formation, VIC activation, microtearing, and compaction in osteogenic-treated groups

We used thrombin (1U/mL) to activate the RhoA pathway in control and osteogenic treated gels, and inhibitor Y-27632 (10uM) to inhibit the RhoA pathway through downstream mediator ROCK[46,47]. We additionally used Rac1 inhibitor NSC23766 (10µM) to inhibit the Rac1 pathway[31]. We previously determined that the homeostatic breakdown in the osteogenic-treated samples occurred after day 4, so we chose to live-track the samples only from day 4 to day 7 (Figure 1*, bottom)*.

The addition of RhoA-activator thrombin and Rac-1 inhibitor to OGM treatment expedited the onset of pathogenic compaction (Figure 7 *A)*[48]. On day 4, OGM+Thrombin and OGM+ Rac1, but not OGM-only treatment groups were significantly more compacted compared to CTRL samples (Figure 7 *A)*. By day 5, OGM, OGM+Thrombin, and OGM+Rac1 groups were all significantly more compacted compared to CTRL, and these groups continued to undergo pathogenic compaction through day 7 (Figure 7 *A)*. Addition of ROCK inhibitor to OGM-treated samples delayed the onset of pathogenic compaction until the last day of culture, at which point some samples began to compact (Figure 7 *A)*. Addition of thrombin to control gels was not sufficient to induce pathogenic compaction. (Figure 7 *A)*.

**Figure 7.**
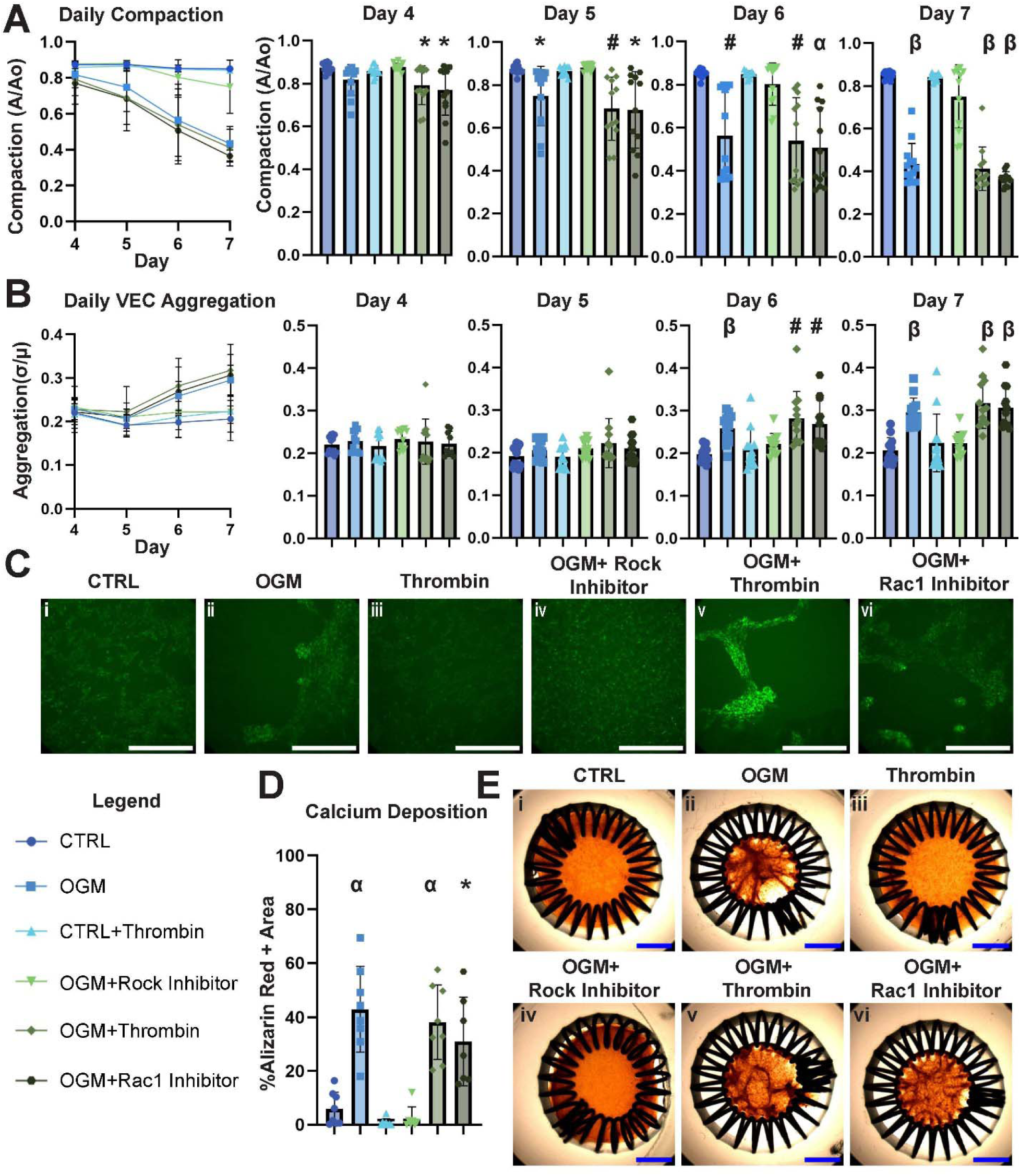
Effect of RhoA-GTPase modulation with thrombin, ROCK inhibitor, and Rac1 inhibitor on compaction, VEC aggregation, and calcification from day 4 to day 7 of culture. A: Compaction plotted over time by treatment group (left) and detailed breakdown by day (right) (n=12-13). B: VEC aggregation plotted over time by treatment group (left) and detailed breakdown by day (right) (n=9-14). C: Representative images of GFP-VEC monolayer aggregation at day 7. Scalebar= 1mm. D: Calcium deposition quantified by alizarin red positive area by group at Day 7 (n=7-8). E: Representative images of alizarin red staining and compaction for each treatment group at day 7. Scalebar= 2mm. One-way ANOVA was performed with Tukey multiple comparisons for all groups compared to control. Brown-Forsythe and Welch ANOVA test with Dunnet’s T3 multiple comparisons was performed on groups with unequal standard deviation. * p≤0.05, # p≤0.01, α p≤0.001, β p≤0.0001

Next, we assessed the role of the RhoA pathway in endothelial delamination and aggregation (Figure 7 *B, C)*. OGM, OGM+Thrombin, and OGM+Rac1 Inhibitor all exhibited significant VEC delamination starting on day 6 and increasing through day 7 compared to CTRL (Figure 7 *B, C)*. Interestingly, OGM+ ROCK inhibitor prevented VEC delamination and aggregation throughout the culture period (Figure 7 *B, C)*. In fact, the monolayer of the VEC cells treated with ROCK inhibitor were more confluent than the CTRL (Figure 7 *B)*.

We performed alizarin red staining to assess the effect of each drug treatment on calcification, and the results are shown in Figure 7D and 7E. OGM, OGM+Thrombin, and OGM+Rac1 inhibitor exhibited large calcific lesions with significantly more calcified area compared to CTRL. OGM+ROCK Inhibitor prevented the formation of calcific lesions, while addition of thrombin to CTRL was not enough to induce the formation of calcific lesions.

We performed combined OCM, confocal imaging on day 5 and day 7 of the GTPase-treated samples (Figure 8 A-E). On day 5, we noticed differences in the VIC morphology, where CTRL and OGM+ROCK inhibitor had much thinner VIC compared to the other groups (day 5, Interior column). Additionally, there was already a slight difference on day 5 in the VEC coverage and morphology, where OGM+Rac1 inhibitor showed some indications of VEC delamination, and the GFP-VEC in control were slightly smaller in size. The endothelial coverage was maintained the most in the OGM+ROCK inhibitor group. There were few obvious differences in the surface tissue between samples on day 5 (surface column). By day 7, the difference in VIC morphology was more apparent, with thicker VIC in the OGM, OGM+Thrombin, and OGM+Rac1 inhibitor groups compared to CTRL and OGM+ROCK inhibitor groups (interior column). Additionally, on day 7 the surface tissue was uniform in the CTRL and OGM+ROCK inhibitor groups compared to the other groups, where large raised features surrounded by empty space were readily observed.

**Figure 8.**
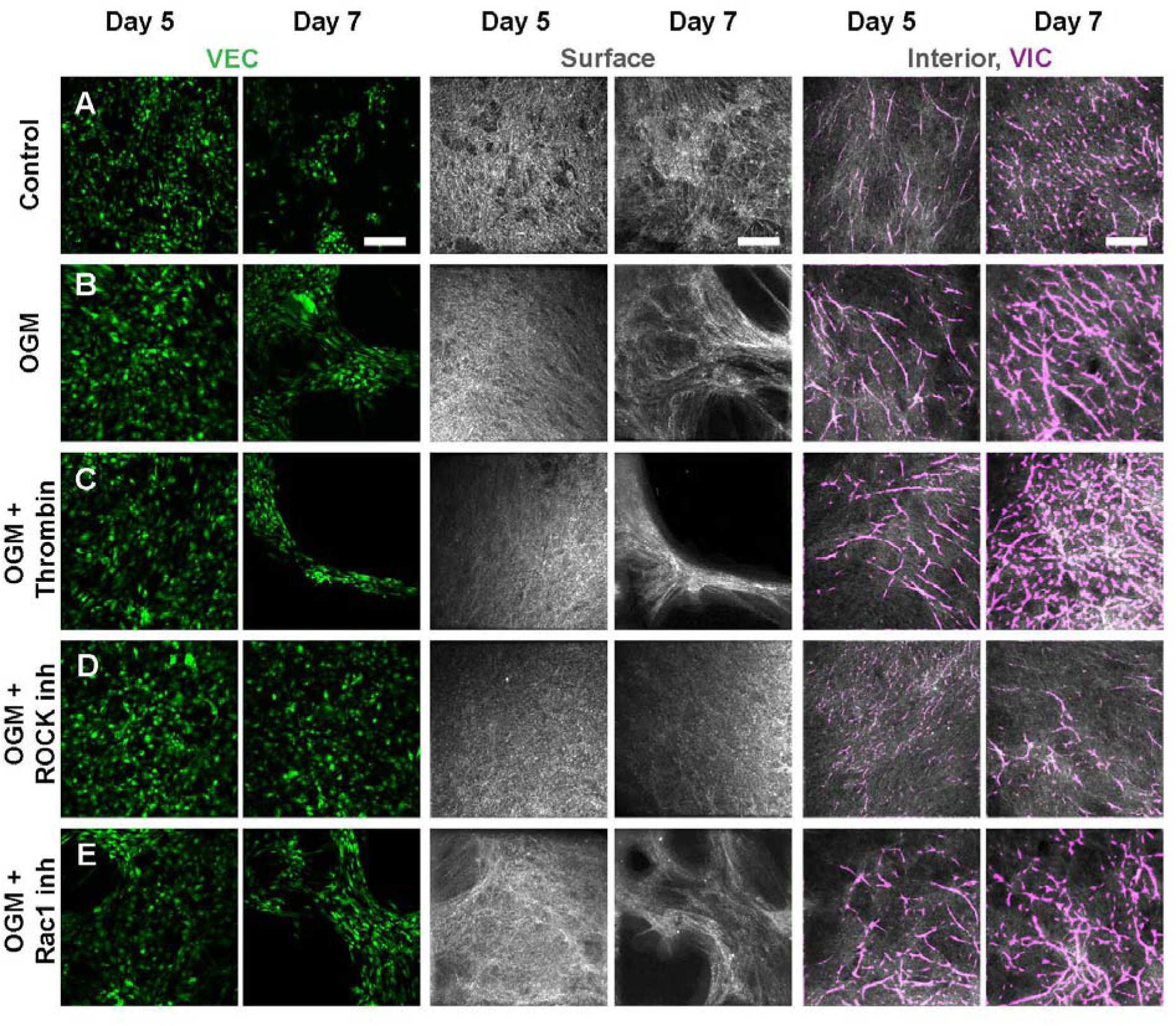
Comparison of pharmacological intervention samples on day 5 and day 7 and graphical summary. A: VEC confocal fluorescence MIP images, OCM MIP at the surface and OCM MIP of a ∼ 20 μm thick section ∼100 μm below the surface with segmented VIC in the magenta channel for the control sample. B. Similar images as part A for OGM samples. C. Similar images as part A for OGM+Thrombin. D. Similar images as part A for OGM+ROCK inhibitor. E. Similar images as part A for OGM+Rac1 inhibitor. Scale bars are 200 μm.

## Discussion and Conclusion

The goal of this study was to identify in real-time the key biological signatures of pathogenic homeostatic breakdown in VEC, VIC, and matrix during calcific lesion formation. To accomplish this, we developed a novel integrated 2D and 3D live imaging platform with GFP-cellular lineage tracing, whereby hourly or daily measurements can be taken to watch the formation of calcific lesions and identify individual cellular contributions to pathogenesis. Using this system, we established that during the formation of raised surface calcific lesions, the VEC monolayer and immediate underlying tissue and interstitial cells spontaneously pull apart and simultaneously aggregate, forming “hills” of raised lesions with VEC aggregation and “valleys” of delaminated endothelium. We also determined that the regions of VEC aggregations directly overlay the regions of calcification *(Graphical Abstract)*.

Injury, inflammation, and dysfunction of the endothelial monolayer have long been postulated as initiating events in CAVD, however these inferences have been drawn largely from the endothelium of already calcified explanted valves[49–53]. A prevailing hypothesis is that the endothelium above regions of calcifications becomes dysfunctional due to a variety of factors including exposure to toxins, hypertension, and mechanical stress, resulting in VIC activation and osteoblastic transformation[54–56]. In this work, we identify for the first time active endothelial delamination/ cell migration mechanism, whereby endothelial monolayer breakdown occurs due to a large-scale “ripping apart” and “clumping” of the endothelium in concert with the underlying interstitium. The regions of endothelial aggregates are the same regions with calcific lesions, whereas the regions immediately surrounding the lesions are delaminated of endothelial cells. A recent investigation of explanted whole-mount human aortic valve leaflets by Morvan et al. identified a unique spatial patterning between regions of calcification, hematomas, and endothelial denudation that are consistent with our findings[57]. In their study, they demonstrated that there is increased iron deposition in correlation to disease severity in calcified valves, indicative of endothelial denudation and subsequent red blood cell (RBC) infiltration into the tissue, and that RBC infiltration likely preceded calcification in valves of varying disease stages. However, they were unable to identify in real time how the endothelial monolayer becomes disrupted leading to downstream coagulation-driven pathophysiology.[57,58]. Our findings in this work provide a plausible mechanism explaining this phenomenon.

Interestingly, the cellular and matrix pathogenesis leading to these large, multicellular lesions at the tissue surface presented separately from other disease processes occurring through the bulk of the tissue. The use of the 1300nm OCM imaging approach enabled much deeper imaging than many standard 3D imaging methods. In particular, the entire sample thickness of the OGM day 7 sample may be imaged as shown in Figure 3D, yielding a complete picture of tissue fibrotic remodeling. Using OCM imaging combined with cross-sectional staining, we identified that deeper regions in our CAVD model exhibited thick, activated αSMA+ myofibroblastic interstitial cells arranged in a “lattice” network with no apparent similarity in pattern to the surface lesions. Tearing of the matrix also occurred in the bulk of the matrix, but presented as small microtears that slowly propagated in contrast to the large-scale delamination that occurred at the surface. Further, in cross-sectional staining, it was clear that the myofibroblasts in tissue depth were more sparsely dispersed compared to the dense-lesion aggregates at the surface.

We originally hypothesized that there might be some correlation between bulk fibrotic matrix remodeling by myofibroblasts, measured by overall tissue compaction, and the delamination/aggregation process leading to lesion formation. Although we did see a weak correlation between the two, we also observed examples of samples with low compaction but high aggregation, indicating that increasing bulk fibrotic remodeling is not necessary to induce increased VEC aggregation and calcific lesion formation. This finding is also consistent with observations in the literature, where some stenotic patients valve explants exhibit extensive fibrotic remodeling without calcification[28,30,59,60]. From these insights, we hypothesize that there might be one initiating mechanism driving both bulk fibrotic remodeling and surface lesion formation that gives way to divergent secondary mechanisms separately driving fibrosis and lesion formation, however further investigation is warranted to determine whether this is true.

Although this is the only study to our knowledge demonstrating a lesion-forming VEC monolayer delamination and aggregation process in CAVD, there some is evidence in non-valve biological systems of pathogenic endothelial monolayer hole formation. Stroka and Aranda-Espinoza found that in the presence of a stiff substrate, exposure to neutrophils caused vascular endothelium to actively form large pathogenic holes for extravasation that could not be closed after the neutrophils had passed through the endothelial monolayer[61]. This aberrant hole formation was due to activation of the Rho-GTPase family, specifically through RhoA and downstream ROCK1/myosin light-chain (MLC)[61]. Huynh et al. also found evidence that the RhoA-GTPase family was responsible for endothelial monolayer breakdown in bovine aortic endothelial cells on stiff substrates[62].

In contrast to the relatively understudied valvular endothelial monolayer, there is extensive evidence in valvular interstitial cells that the RhoA, and to a lesser extent Rac1 pathways play a role in CAVD. RhoA/ROCK has been shown to be activated in VIC by a number of calcific-nodule inducing stimuli, including oxLDL, LPA, OGM, Cadherin-11 overexpression, fibrin, mechanical strain, and inorganic phosphate medium[32–38]. RhoA/RhoA-GTP elevation has also been demonstrated in calcified human aortic valve explants[32,33]. Similarly, active Rac-1 has been found to be elevated in calcified human aortic valves, and its inhibition in VIC cells alone and free-floating porcine valve explants with NSC23766 was able to reduce OGM-induced calcification[31].

In this work, immunofluorescent cross-sectional staining of OGM-samples revealed an abundance of both Rho-A GTP and Rac-1 GTP positive cells, although the Rac1-GTP positive cells were found in both OGM and control groups while the RhoA-GTP positive cells were only found in the OGM samples. Our GFP-VEC lineage tracing approach was able to establish that this Rho-A GTP upregulation was present in both the VIC and VEC in osteogenic-treated samples. Pharmacological inhibition of Rho-A with ROCK inhibitor Y27632 prevented multiple aspects of pathogenesis including VEC aggregation, compaction, and calcium deposition. There was no delamination and tearing of the endothelial basement membrane observed with ROCK inhibition, similarly to control samples. Further, there was no evidence of VIC thickening or deep-microtear formation often seen in osteogenic-treated samples. Conversely, activation of RhoA GTP signaling in OGM-samples with Thrombin was sufficient to induce earlier bulk tissue compaction, however, activation of Rho-A GTP with Thrombin alone was not enough to induce a significant increase in compaction, aggregation, or calcium deposited-area in control samples. In sum, this indicates that Rho-A GTP signaling contributes substantially but not wholly to the CAVD pathogenesis observed in our biomaterial-model.

In contrast, inhibition of Rac-1-GTP was ineffective in preventing VEC aggregation and calcium deposition, and induced an enhanced fibrotic response measured by bulk tissue compaction, similarly to the OGM+Thrombin treatment group. This effect of Rac-1 inhibition is in contrast to our previous findings with VIC-only and porcine CAVD models, indicating a likely differential cell-specific role of Rac-1 in CAVD pathogenesis and emphasizing the need for models of CAVD to capture both VIC and VEC to fully faithfully recapitulate CAVD pathogenesis[31]. There is some evidence of a mutually inhibitory relationship between Rac-1 GTP and Rho-A GTP, which could help explain why inhibition of Rac-1 induced a similar pathogenic response to Rho-A activation with thrombin, although further investigation is warranted to confirm whether this mechanism explains the disease-promoting effects of Rac-1 inhibition we observed[26,46,63,64]. Because RhoA and Rac1 are usually mutually inhibitory, it is unusual to see both pathways upregulated at the same time, as was seen in the immunofluorescent staining of our osteogenic treated samples. Interestingly, in glioblastoma cells, forced double activation of RhoA and Rac1 resulted in an emergent aggregation behavior that was not seen in RhoA or Rac1 only activation[65].

Work by Arevalos et al. also investigated the role of RhoA/Rac1 in porcine valvular endothelial cells, but through a vasculogenic lens[66]. They found that ROCK inhibitor Y27632 decreased cell migration and lacunarity (empty space between neovessels), while Rac-1 inhibitor NSC23766 had no effect on cell migration, and increased VEC lacunarity[67]. Additionally, they determined with immunostaining that ROCK-inhibited samples exhibited a flatter morphology with a looser cell network, whereas Rac-1 inhibition resulted in dense assemblies of endothelial cells[67]. These observations are in line with our VEC aggregation and lesion formation results, where ROCK inhibitor prevented VEC migration and 3D aggregation, while RAC-1 inhibition had no effect on this process. In previous experiments (unpublished), we have not observed that GFP-VEC seeded in a monolayer have the propensity to form vascular networks on collagen-only hydrogels treated with osteogenic medium. However, there are some distinct similarities in the patterning of the delaminated areas of endothelium in the lesion space of our osteogenic-treated VIC-VEC co-culture samples and the lacunae seen in the monoculture tube-formation assay[67]. Investigation on whether there is any additional mechanistic overlap in the formation of vessel networks/lacunae in vasculogenesis and the formation of VEC monolayer aggregations leading to calcific nodules in CAVD could prove promising for future investigation.

The rapid onset and extent of endothelial delamination and clumping revealed by this work may be alternatively caused by tensile forces present across the entire endothelial monolayer that are locally disrupted due to a breakdown in cell-cell junctions resulting in a “recoil effect” that causes the endothelial cells to aggregate in some regions and cleave in others. Laser ablation to intact embryonic epithelial monolayers induce similar local epithelial fenestrations indicative of residual tensile stress, and could be a promising avenue for further investigation[68,69]. Another plausible instigator of VEC delamination could be local migration, proliferation, or contraction of RhoA-GTPase activated VIC directly underlying the VEC monolayer. This phenomenon could induce an expansive pressure from below, causing the endothelium directly on top of these VIC bundles to rip apart from the surrounding monolayer.

Now that the onset of lesion formation has been established in this work with live GFP-VEC, VIC, and matrix imaging, evaluation of the efficacy of therapeutic administration at different stages and aspects of CAVD pathogenesis is possible. We have demonstrated that onset of 3D surface lesion formation may be readily observed with live GFP-VEC OCM/Confocal imaging, allowing future work to investigate activation of different molecular mechanisms and efficacy of promising therapeutics before, during, or after the onset of lesion formation. Additionally, we have delineated differential disease processes that occur at the VEC-VIC interface vs. deeper in the bulk of the tissue, allowing for distinction of which pathogenic processes future therapeutic targets of interest act upon. This opens opportunities for more customizable patient-specific therapeutic evaluation.

Altogether, from our approach and findings in this work, we (1) identified a new temporal pathophysiological mechanism of homeostatic breakdown in the valvular endothelium in the onset of CAVD, (2) demonstrated the utility of a new multimodal imaging workflow for CAVD investigation, in which both VIC and VEC cells can be independently traced live in a physiologically relevant 3D biomaterial construct, and (3) utilized this imaging approach with the pathophysiological biomarkers we established in control and osteogenic-treated samples to evaluate whether and how modulation of the RhoA and Rac1 pathways might prevent these disease processes. This new platform in conjunction with the new knowledge of pathophysiology gained from this study will allow for more nuanced and spatiotemporally-relevant evaluation of therapeutics for efficacy in preventing and treating CAVD.

## Acknowledgements

We thank Timberline Meats in Dundee, NY for their generous donation of porcine hearts. We would also like to thank the Cornell Institute of Biotechnology (RRID: SCR_021741, RRID:SCR_021740) for their assistance with confocal microscopy and flow cytometry, particularly Dr. Johanna Dela-Cruz. We would like to thank Dr. Warren R. Zipfel for his inputs and for loaning us the confocal laser and photomultiplier tubes. We would like to acknowledge Dr. Justin Luo and Dr. Meg McFetridge with their assistance in optimizing the optical coherence microscopy imaging technique. We would also like to acknowledge Dr. Terence Gee and Dr. Lara Estroff for their mentorship. This work was sponsored by the National Institute of Health (F31HL165886, HL128745, HL143247, HL160028, R01GM132823, R21CA253197). Graphics in Figure 1 and Graphical Abstract were Created with BioRender.com.

## Data availability

We are unable to share the data at this time due to its size, complexity, and custom file extensions

## Glossary

**Abbreviation**
2D: Two-Dimensional
3D: Three-Dimensional
aSMA: Alpha-Smooth Muscle Actin
AZR: Alizarin Red
BSA: Bovine Serum Albumin
CAVD: Calcific Aortic Valve Disease
CDH5: VE-cadherin
CTRL: Control Medium
DMEM: Dulbecco’s Modified Eagle Medium
EndMT: Endothelial to Mesenchymal Transition
FACS: Fluorescence Activated Cell Sorting
FBS: Fetal Bovine Serum
FOV: Field of View
GFP: Green Fluorescent Protein
IF: Immunofluorescence
LPA: Lipoprotein(a)
MIP: Maximum Intensity Projection
MLC: Myosin Light Chain
OCM: Optical Coherence Microscopy
OGM: Osteogenic Medium
oxLDL: Oxidized Low-Density Lipoprotein
P/S: Penicillin Streptomycin
PBS: Phosphate Buffered Saline
PBST: Phosphate Buffered Saline + Tween-20
PDMS: Polydimethylsiloxane
PECAM-1: Platelet Endothelial Cell Adhesion Molecule 1
PFA: Paraformaldehyde
Rac-1: Rac Family Small GTPase 1
RBC: Red Blood Cell
RhoA: Ras Homolog Family Member A
RhoB: Ras Homolog Family Member B
RhoC: Ras Homolog Family Member C
ROCK: Rho Associated Coiled-Coil Containing Protein Kinase 1
VEC: Valvular Endothelial Cells
VIC: Valvular Interstitial Cells

