## Supplementary Figure Captions for "Valve endothelial monolayer fissuring via RhoA activity induces 3D calcific aortic valve lesion emergence as revealed by a longitudinal live-imaging platform"

**Supplementary Figure S1. GFP-VEC transfection validation and daily image intensity in control and OGM-treated samples.** A: Brightfield image of GFP+ VEC morphology. B: Fluorescence image of GFP-VEC (green) and nuclei (DAPI). C: GFP-Transfection effeciency quantified by primary cell lot (n=3-6). D-E: 2^-ddCT Fold-change aSMA and PECAM1 mRNA expression for WT-VIC, WT-VEC, and GFP-VEC normalized to WT-VEC (n=4-8). F: Daily average pixel intensity summary of CTRL and OGM GFP-VEC normalized by day (n=34-35). G: Daily breakdown of data summarized in (F). Scalebars=200µM. Student’s t-test performed for (F) and (G). Welch’s T-test was performed in cases of unequal variance. *** p≤0.001, **** p≤0.0001.

**Supplementary Figure S2. Detailed breakdown of aggregation, compaction, and calcium endothelial overlay analysis.** A: Breakdown of daily GFP-VEC aggregation by day (n=34-36). B: Breakdown of daily compaction by day(n=38-44). C: Example workflow of GFP/AZR overlay segmentation and quantification. D: Percent AZR+ GFP+ area normalized to AZR+ GFP- area (n=14).Student’s t-test was performed on control vs. OGM samples in A-B, and AZR+GFP+ vs AZR+ GFP- samples in D. Welch’s T-test was performed in cases of unequal variance. **** p ≤ 0.0001 Figure 3. Comparison of the same control and OGM samples on Day 2 and Day 7.

**Supplementary Figure S3. Cell Segmentation and analysis.** A. Day 7 Control sample VIC analysis, (i) OCM below surface MIP, (II) VIC cell segmentation, (iii) Identified primary objects used for measurements. B. Day 7 OGM sample VIC analysis, (i) OCM below surface MIP, (II) VIC cell segmentation, (iii) Identified primary objects. C. Day 2 OGM sample VEC analysis (i) GFP fluorescence MIP, (ii) Identified primary objects used for measurements. D. Day 7 OGM sample VEC analysis, (i) GFP fluorescence MIP, (ii) Identified primary objects. E. Validation of VIC segementation with a GFP tagged VIC sample, (i) OCM below surface MIP, (II) VIC cell segmentation, (iii) Identified primary objects from the segementation used for measurements, (iv) VIC fluorescence MIP, (v) Identified primary objects from the Confocal fluorescence MIP used for measurements. F. Comparison of measurements made on (iv) and (ii). G. VIC morphology assessed on Day 7 using Phalloidin staining and Zeiss LSM 880 on (i) Control, (ii) OGM sample. Scale bars are 200 µm.
