## Supplementary figures and images for "Valve endothelial monolayer fissuring via RhoA activity induces 3D calcific aortic valve lesion emergence as revealed by a longitudinal live-imaging platform"

### Supplementary Figure S1

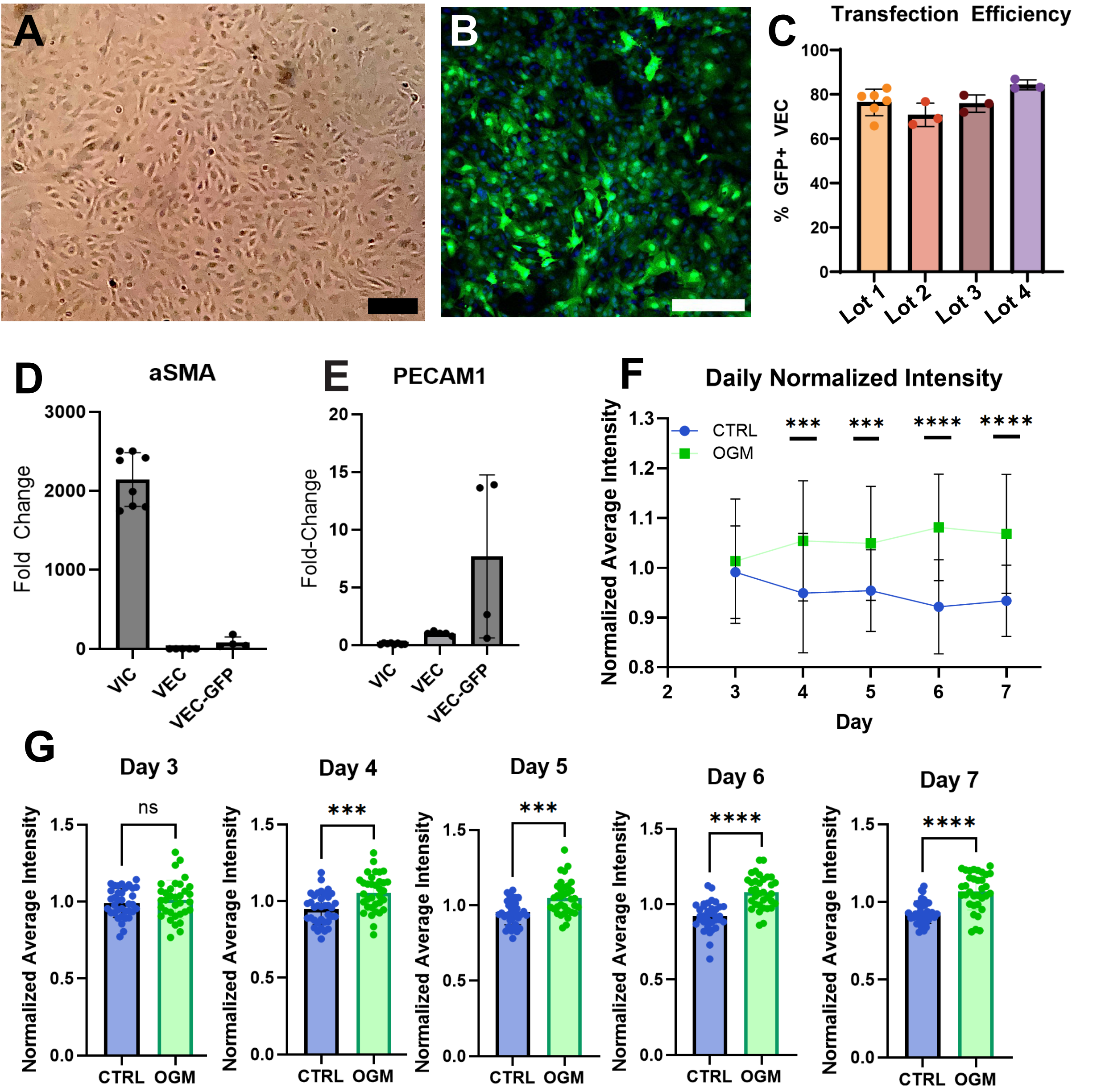

### Supplementary Figure S2

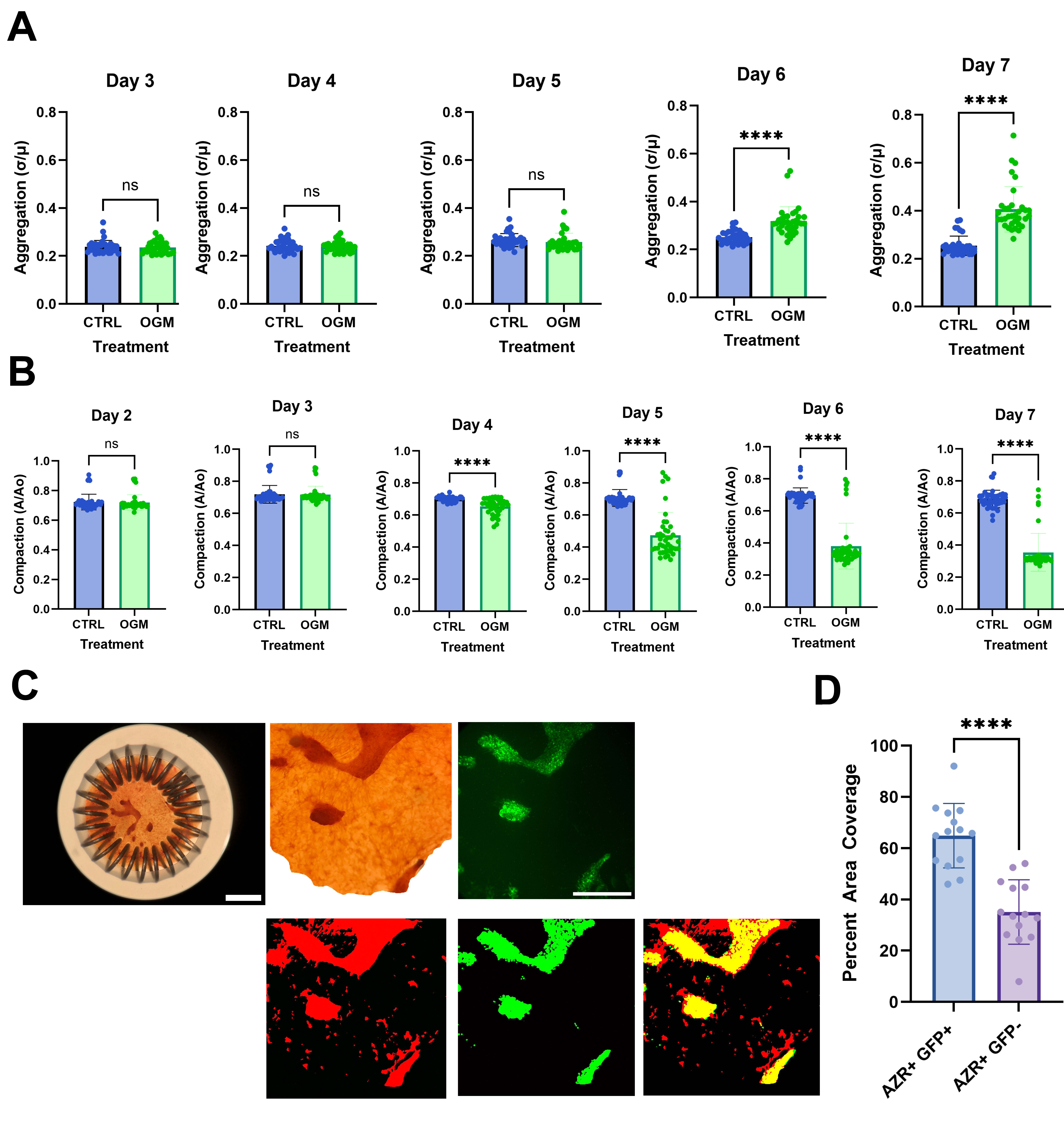
