## Supplementary Figure S3 for "Valve endothelial monolayer fissuring via RhoA activity induces 3D calcific aortic valve lesion emergence as revealed by a longitudinal live-imaging platform"

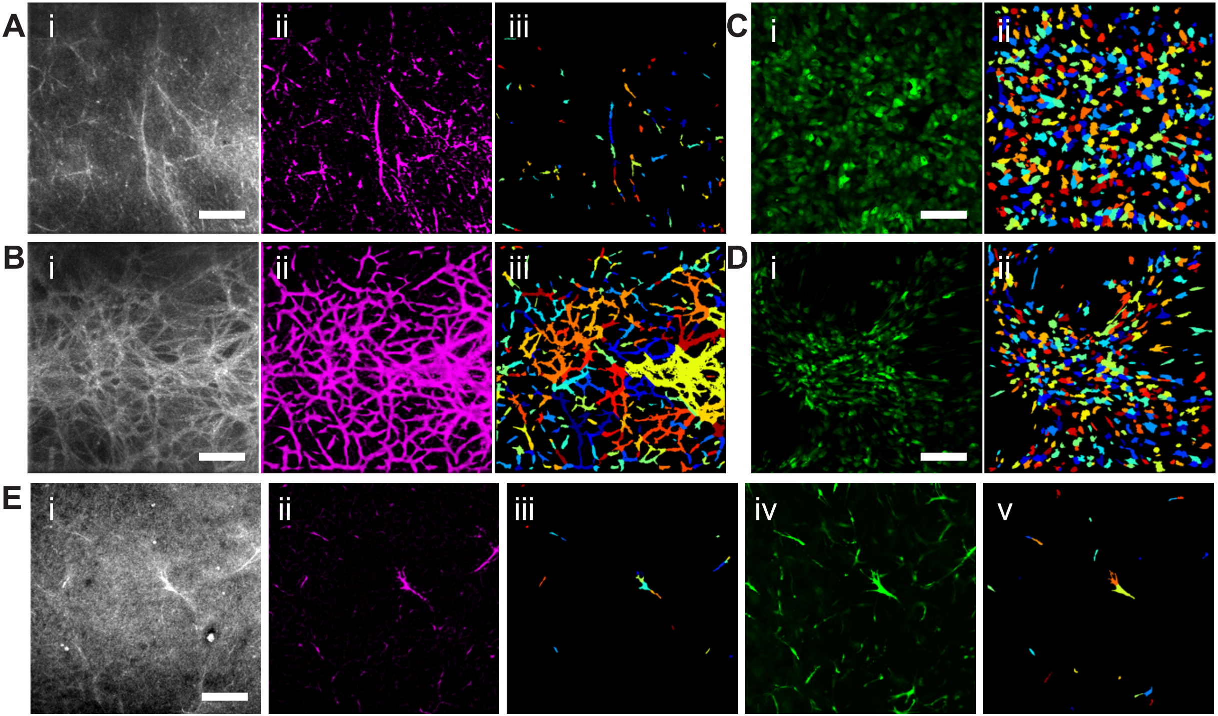


F

| Measurement | Confocal mean | Confocal stdDev | OCM mean | OCM stdDev |
|---|---|---|---|---|
| Cell Count | 21 | | 19 | |
| Cell Area Occupied (%) | 0.70 | | 0.41 | |
| Cell Area (µm²) | 333.33 | 343.03 | 214.93 | 191.13 |
| Cell mean radius (µm) | 2.25 | 0.63 | 2.06 | 0.56 |
| Cell Eccentricity | 0.92 | 0.08 | 0.88 | 0.13 |
| Cell minFeretDiam (µm) | 13.19 | 8.54 | 10.55 | 7.12 |
| Cell maxFeretDiam (um) | 38.43 | 21.62 | 28.29 | 14.10 |

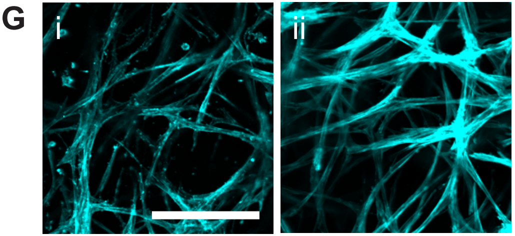
